# Natural dispersal is better than translocation for reducing risks of inbreeding depression in eastern black rhinoceros (*Diceros bicornis michaeli)*

**DOI:** 10.1101/2023.06.28.546899

**Authors:** Ronald V. K. Mellya, J. Grant C. Hopcraft, William Mwakilema, Ernest M. Eblate, Simon Mduma, Bakari Mnaya, Idrissa Chuma, Emmanuel S. Macha, Dickson Wambura, Robert Fyumagwa, Elizabeth Kilbride, Barbara K. Mable, Anubhab Khan

**Affiliations:** School of Biodiversity, One Health and Veterinary Medicine, University of Glasgow, Glasgow, G12 8QQ, United Kingdom; Tanzania National Parks (TANAPA), P.O. Box 3134, Arusha, Tanzania; Tanzania Wildlife Research Institute (TAWIRI), P.O. Box 661, Arusha, Tanzania; Serengeti Biodiversity Programme, P.O. Box 661, Arusha, Tanzania; Ngorongoro Conservation Area Authority (NCA), P.O. Box 1, Ngorongoro, Arusha, Tanzania; Wildlife Conservation Initiative (WCI), P. O. Box 16020, Arusha, Tanzanial

**Keywords:** conservation management, whole genome sequencing, endangered species, inbreeding, genetic load

## Abstract

Due to ever increasing anthropogenic impacts, many species survive only in small and isolated populations. Active conservation management to reduce extinction risk includes: increasing habitat connectivity; translocations from captive populations; or intense surveillance of highly protected closed populations. The fitness of individuals born under these scenarios may vary due to differences in selection pressures. However, the genetic impacts of such strategies are rarely assessed. Using whole genome sequences from cohorts of the critically endangered eastern black rhinoceros as a model, we compare the consequences of past conservation efforts. We find that offspring of individuals that had either dispersed from native populations (F_ROH>100Kb_ = 0.13) or translocated from captive populations (F_ROH>100Kb_ = 0.08) showed lower inbreeding compared to closed populations (F_ROH>100Kb_ = 0.17). However, the frequency of highly deleterious mutations was higher for offspring resulting from translocation compared to the other groups and this load was sheltered by higher heterozygosity. This could increase risks of inbreeding depression if captive founders subsequently inbreed after translocation. In contrast, native dispersers reduced the negative effects of inbreeding without compromising the benefits of past purging of deleterious mutations. Our study highlights the importance of natural dispersal and reiterates the importance of maintaining habitat corridors between populations.

## Introduction

Many highly threatened animal species persist in small, isolated patches that are susceptible to inbreeding and loss of genetic variation due to drastic reductions in population size (i.e. bottlenecks or founder effects), which are warning signals of populations at threat of extinction (Hoban et al. 2022). When too few individuals remain in the wild or when there is insufficient genetic variation for natural dispersal to have a substantial impact, active human interventions have been used for genetic rescue (Weeks et al. 2011; Frankham 2015; Seddon & Schultz 2020). This includes translocation of individuals between native populations (*in situ*) or (re)-introduction from captive populations (*ex situ)* (Berger-Tal et al. 2020). However, such assisted movement has often been conducted without explicitly assessing existing patterns of genetic variation or basing decisions on only a handful of genetic markers (Mable 2018; Purisotayo et al. 2019), which has not allowed assessment of the long-term fitness impacts of past or future management decisions (Ralls et al. 2018; Ralls et al. 2020).

Whole genome sequencing data has the potential to revolutionise genetic rescue because of the expanded inferences possible compared to single-gene approaches. Detailed cost-benefit analyses can be conducted not only to select which individuals might be most beneficial to move but also to predict the genome-wide fitness consequences of different intervention scenarios (Robinson, J. A. et al. 2019; Saremi et al. 2019; Hoffmann et al. 2021; Khan et al. 2021). Additionally, inferences about demographic history can be modelled more accurately based on millions of single nucleotide polymorphism (SNP) loci compared to markers like microsatellites. This is important to enable assessment of the potential impacts of previous bottlenecks to purge deleterious recessive mutations (Frankham et al. 2001; Hedrick & Garcia-Dorado 2016; Grossen et al. 2020; Khan et al. 2021). However, artificial management strategies designed to boost wild populations such as establishing founder sub-populations or reintroducing captive animals into the wild could inadvertently negate the long-term benefits of purging deleterious alleles in surviving populations. The issue of primary concern is that the genomic consequences of different management strategies in relation to inbreeding and deleterious mutation loads has rarely been assessed. Here we use critically endangered eastern black rhinoceros (*Diceros bicornis michaeli*) populations in Tanzania as a model for predicting the relative fitness impacts of *ex situ* conservation, translocations from captive populations to the wild, and natural dispersal through corridors to gene flow. The species is critically endangered due to extensive poaching across their natural range, with most surviving individuals restricted to highly protected areas such as zoos, closed sanctuaries and intensive protection zones (IPZ) (Emslie et al. 2009; Fyumagwa & Nyahongo 2010; Muya et al. 2011; Moodley et al. 2017). Active management has so far succeeded in preventing their extinction in the short term, but previous translocations have not been informed by genetics (Mellya et al. 2023) and long-term consequences remain unknown.

Specifically, we use whole genome sequencing data to assess: 1) the scale of inbreeding that has been induced by the severe bottlenecks and subsequent expansion of native populations; and 2) what impacts previous attempts at population supplementation have had on the accumulation of potentially deleterious mutations. Our main aim is to question the potential trade-offs between: 1) increasing adaptive potential by introducing new variation into threatened populations; and 2) increasing risks of inbreeding depression (reduction in fitness due to inbreeding) caused by introducing deleterious mutations (genetic load). We hypothesis that the genetic load might have been purged from inbred wild populations (due to severe past bottlenecks and ongoing inbreeding) but “hidden” in captive populations (due to reduced selection under the benign conditions under which they are kept or increased heterozygosity due to mixing individuals from different sources). We consider the impacts of various management practices on: (i) genetic diversity; (ii) genome-wide inbreeding; (iii) accumulation of deleterious mutational load (relative load); and (iv) homozygosity of deleterious alleles (realised load). We also assess the potential for purging by using the whole genome sequence data from native individuals to estimate the timing and extent of previous bottlenecks. We provide recommendations for managing eastern black rhinoceros populations, as well as more general strategies for genetic rescue of other highly endangered species.

## Results

### Observational pedigrees

Based on observational records and pedigrees (Supplementary Figures 1-4), we divided the extant Serengeti-Mara subpopulations into five cohorts (Figure 1a; Table 1; Supplementary Table 1), defined as: 1) native, no dispersal (NN) - individuals with two wild parents from the same native subpopulation; 2) recent natural dispersal (rND), - first generation offspring of individuals who dispersed from a native subpopulation and mated with an individual in their new resident subpopulation; 3) old natural dispersal (oND) – 2^nd^ or 3^rd^ generation offspring of individuals that had dispersed between native subpopulations but where there has been no subsequent gene flow into that lineage; 4) translocation/assisted dispersal (AD) - individuals where one parent was native to the population and the other was born in a captive breeding facility; and 5) translocated captive (TC) - individuals that were translocated to Tanzania but whose parents were both born in captivity.

**Figure 1:**
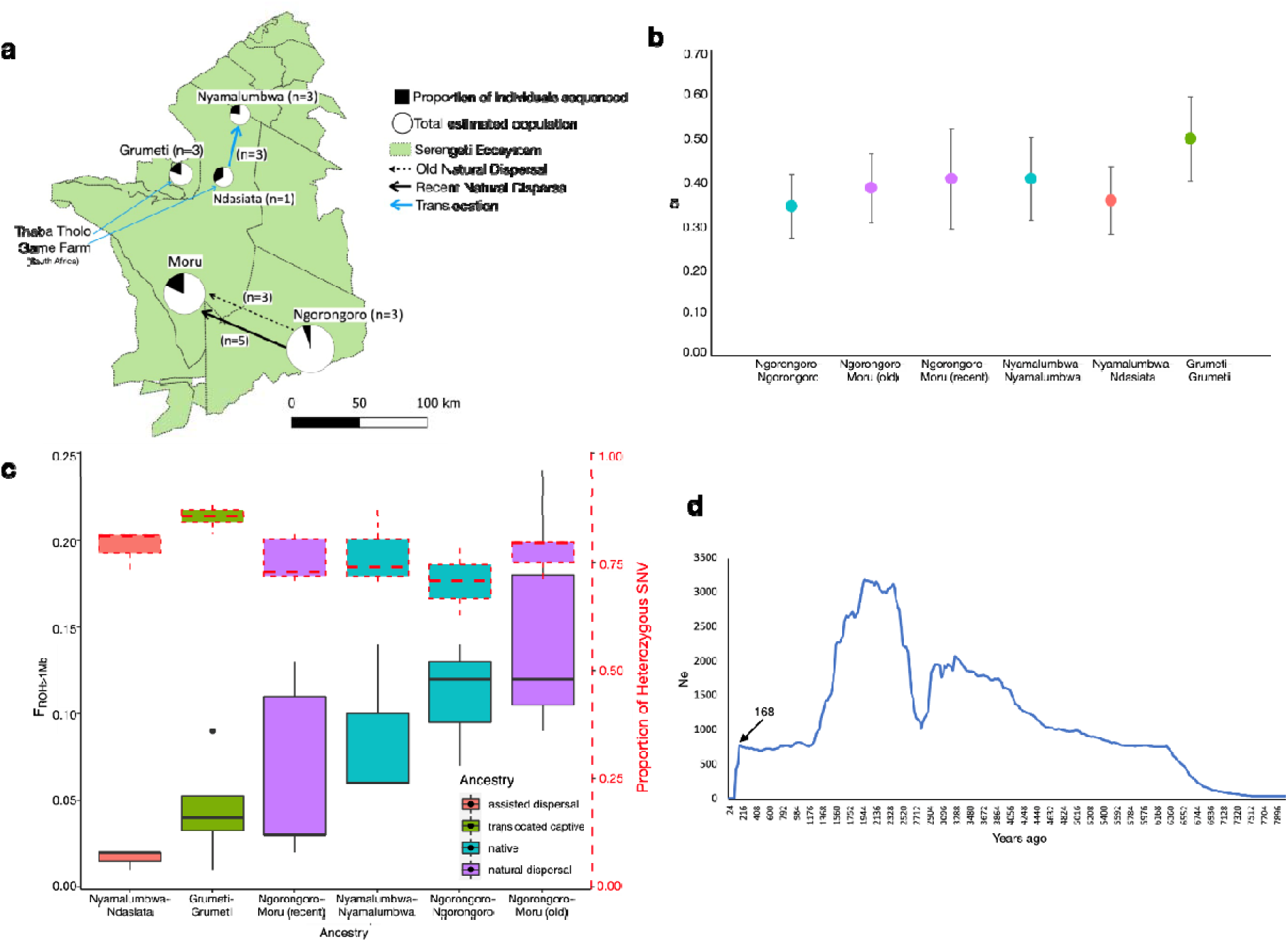
Sampling strategy, impact of ancestry of cohorts and demographic history. (a) Map of the Serengeti ecosystem showing the sampling locations, with the size of the pie chart being proportional to the total estimated population size and black being proportional to the number of offspring within the population that were sequenced. Old natural dispersal are 2^nd^ and 3^rd^ generation offspring of individuals that dispersed naturally whereas recent dispersal refers to movement of the parents of the individual sequenced. The impact of different translocations on: (b) genetic diversity, measured as the average number of pairwise differences between individuals in a cohort (pi); (c) inbreeding, measured as ROH stretches of 1Mb, indicative of recent inbreeding, with boxplots outlined in red dotted lines indicating proportion of heterozygous single nucleotide variants (SNV) in polymorphism-containing loci; and (d) the recent demographic history of eastern black rhinoceros populations, with the arrow indicating the time of the most recent bottleneck.

**Table 1:** Ancestry of sampled cohorts, indicating the source of the parents, the type of ancestry (NN = native, no dispersal; rND = recent natural dispersal; oND = old natural dispersal; AD = assisted dispersal; TC = translocated individuals whose parents were from captive populations), the subpopulation from which the individual was sampled, and the number of individuals sequenced.

| Sire | Dame | Ancestry type | Subpopulation | N |
| --- | --- | --- | --- | --- |
| Ngorongoro | Ngorongoro | NN | Ngorongoro | 3 |
| Ngorongoro | Moru | rND | Moru | 5 |
| Ngorongoro | Moru | oND | Moru | 3 |
| Nyamalumbwa | Nyamalumbwa | NN | Nyamalumbwa | 3 |
| Nyamalumbwa | Ndasiata | AD | Ndasiata | 3 |
| Captive-Thaba | Captive-Thaba | TC | Grumeti/Ndasiata* | 4 |
| Tholo | Tholo |  |  |  |

\* Since only a single translocated individual was available from Ndasiata, this was combined with the adjacent Grumeti population, which was established from the same captive population in South Africa.

Individuals presently in Moru are the most inbred because the population had only three founders and the pedigree clearly demonstrates transgenerational mating and skew of reproductive effort (Supplementary Figure 1). However, the single founding male (R21) had dispersed naturally from the native Ngorongoro population, meaning that all rND and oND individuals descended from that sire (Table 1; Supplementary Table 1; Supplementary Figure 1). The native Ngorongoro population experienced a milder bottleneck than Moru but is still characterised by multiple generations of inbreeding (Supplementary Figure 2). The records for the other native population in Tanzania (Nyamalumbwa) are not as complete because it is a transborder population connected to Masai Mara National Reserve in Kenya (Supplementary Figure 3), and there has not been a common system of monitoring developed between the two countries. For the Ndasiata subpopulation, founded by South African captive individuals, the samples sequenced were offspring of a native individual who had dispersed from Nyamalumbwa (ND10) and mated with captive founders from Thaba Tholo Game Reserve in the Republic of South Africa; these were classified as assisted dispersal (AD, Supplementary Figure 4). We also had a sample available from one of the captive founders (R9) from Ndasiata in Tanzania, but this was grouped with the translocated captive (TC) individuals from the adjacent Grumeti population (Grumeti Game Reserve) for subsequent analyses because some of the founders were from the same captive population.

### Genetic diversity

Using whole genome sequencing (at least 10x coverage) of 3-5 individuals per cohort, we found similar genetic diversity across cohort types, except for the TC individuals (Grumeti/Ndasiata), which showed higher pairwise nucleotide diversity (pi; Figure 1b) and allelic richness (Supplementary Figure 5) than the others. The Ngorongoro native individuals showed the lowest allelic richness, while we found no significant difference between AD and Nyamalumbwa native individuals. The oND and rND individuals did not differ from one another in allelic richness but both had higher nucleotide diversity than the Ngorongoro native individuals. Pairwise relatedness (Supplementary Figure 6) was highest for the assisted dispersal cohort (Nyamalumbwa-Ndasiata) but lowest for the individuals native to Nyamalumbwa. This was also reflected in the Principal Components Analysis (Supplementary Figure 7), which shows more spread along axis 2 than for the other populations.

### Genome-wide Inbreeding

Based on runs of homozygosity (ROH) longer than 1 Mbp (historical inbreeding; Figure 1c) and more than 20 Mbp (recent inbreeding; Supplementary Table 1; Supplementary Figure 8), both assisted dispersal (Nyamalumbwa-Ndasiata) and recent natural dispersal (Ngorongoro-Moru) showed lower levels of inbreeding compared to individuals with parents that had not moved (Ngorongoro-Ngorongoro, Nyamalumbwa-Nyamalumbwa; Figure 1c). Levels of inbreeding in the native individuals were as predicted by the observational pedigrees, with Nyamalumbwa individuals showing substantially less historical inbreeding than Ngorongoro individuals (Figure 1c). However, the offspring of older dispersal from Ngorongoro to Moru were as inbred as the offspring of native Ngorongoro individuals (Figure 1c; Supplementary Table 1). Interestingly, the assisted dispersal cohort showed even less historical (Figure 1c) and recent inbreeding (Supplementary Table 1) than the captive individuals that had been translocated to Grumeti or Ndasiata.

Based on the proportion of single nucleotide variants (SNVs) that are heterozygous, both recent (rND) and old natural dispersal (oND) were equally effective at increasing heterozygosity compared to individuals that had not dispersed (Figure 1c). However, while assisted dispersal (Nyamalumbwa-Ndasiata) did not substantially alter heterozygosity compared to the native Nyamalumbwa cohort, the cohort involving only translocated individuals (Grumeti/Ndasiata) had the highest average heterozygosity (Figure 1c).

### Recent demographic history

Estimates of recent demographic history using only the individuals with two native parents (NN, rND, oND cohorts) suggest that eastern black rhino populations had an effective population size less than 3500 in the last 350 generations (8400 years) (Figure 1d). Additionally, the population faced a severe bottleneck 80 generations (∼1900 years) ago and then again 7 generations (∼168 years) ago.

### Accumulation of genetic load

Based on derived missense mutations (expected to be mildly deleterious), individuals whose parents had dispersed recently (rND; Ngorongoro-Moru) or individuals whose grandfather (or great-grandfather) had moved naturally from Ngorongoro to Moru (oND) showed higher genetic load than their native source population (NN; Ngorongoro) but showed a similar load compared to the assisted dispersal cohort (AD; Ndasiata-Nyamalumbwa) and to individuals with both parents native to Nyamalumbwa (Figure 2a; Supplementary Figure 9a). The AD cohorts showed a lower load compared to the translocated captive (TC) individuals but both cohorts involving captive parents had higher load compared to Ngorongoro natives (NgNg; Supplementary Figure 9a). Native individuals with both parents from Nyamalumbwa (NyNy) had loads similar to translocated captive (TC) individuals (Supplementary Figure 9a).

**Figure 2:**
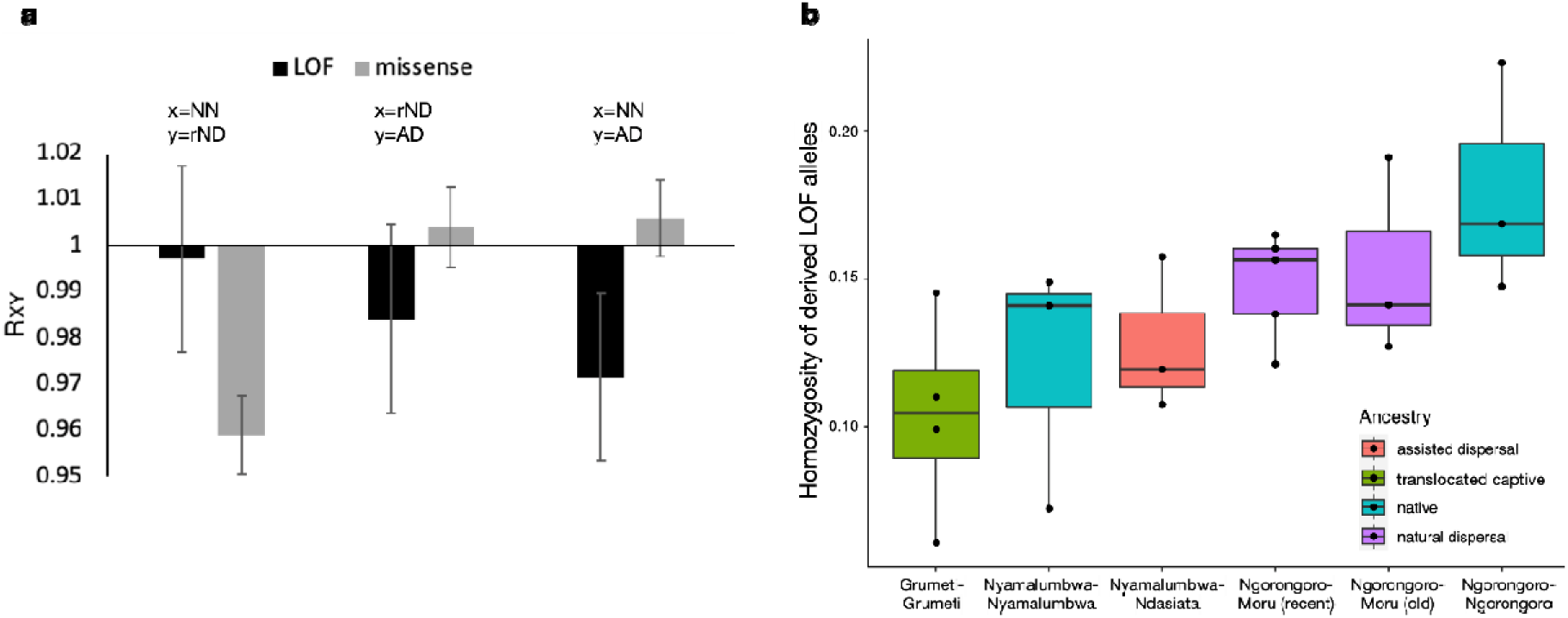
R**e**lative **(a) and realised (b) deleterious allele load.** Relative deleterious allele load, measured as that in the cohort “x “with respect to the cohort “y” listed (see Supplementary Figure 9 for a comparison across all cohorts), is shown for both mildly deleterious (missense) and more serious (loss of function, LOF) mutations. NN represents offspring of parents native to an area, rND represents first generation offspring of a native and dispersed parent, AD represents offspring of a native and a translocated parent. For NN-rND comparisons, individuals of Ngorongoro-Ngorongoro were compared to those with Ngorongoro-Moru ancestry. For NN-AD comparisons, individuals of Nyamalumbwa-Nyamalumbwa were compared to those with Nyamalumbwa-Ndasiata ancestry. (b) Homozygosity of derived LOF alleles in various ancestries.

Based on more detrimental loss of function mutations (LOF), there were no differences between individuals whose father had recently dispersed (rND) and individuals from his native source subpopulation (NN). More substantial differences were observed between the assisted dispersal cohort and both their native population (NN) and the rND cohort (Figure 2a). Strikingly, the old native dispersal cohort (oND) showed evidence of purging of the most serious mutations, as they showed lower LOF loads than any of the other cohorts (Supplementary Figure 9b). Native individuals with both parents from the same location (either Ngorongoro or Nyamalumbwa), did not differ from one another and showed lower loads than cohorts including captive relatives (Supplementary Figure 9b).

Enrichment analysis of the LOF mutations showed a high frequency of genes associated with myosin complex and energy related functions, as well as cytoskeletal functions and anion binding (Supplementary Figure 10), which could suggest relative fitness consequences.

### Realised genetic load

Homozygosity of both derived LOF (Figure 2b) and missense (Supplementary Figure 11) mutations was highest for the native Ngorongoro cohort but both recent and older natural dispersal decreased homozygosity of deleterious alleles. Although assisted dispersal did not result in a change in homozygosity compared to the native Nyamalumbwa cohort, the native Nyamalumbwa cohort had lower homozygosity of deleterious alleles than Ngorongoro. Translocated captive individuals (Grumeti/Ndasiata) showed the lowest homozygosity of deleterious alleles.

## Discussion

Overall, our results question some of the assumptions of *ex situ* conservation strategies: past translocations from captive populations have achieved the goals of increasing the overall population size, as well as the genetic variation and heterozygosity, but at what cost? Introducing new alleles and increased heterozygosity of beneficial alleles could increase adaptive potential but our results also emphasise that introduction of deleterious alleles that have been sheltered in heterozygotes could result in increased inbreeding depression if exposed as homozygotes in the wild. This is consistent with the observation from meta-analyses that translocation of captive-born individuals is often less successful than translocation of wild-born individuals (Griffith et al. 1989; Rummel et al. 2016). The short-term benefits of introducing genetic variation from different sources could thus be compromised in the long term unless there are sustained efforts to reduce subsequent inbreeding (Pérez-Pereira et al. 2022).

Long-distance translocations are expensive not only in terms of financial costs and logistics but also can come at a cost to animal welfare (Teixeira et al. 2007). Encouragingly, at least for highly threatened eastern black rhinoceros, we find that both assisted and natural dispersal reduce inbreeding in the target population. However, natural movement of individuals between subpopulations results in a lower high-impact mutational load of deleterious alleles than mating between wild and translocated individuals. This could suggest that purging due to past bottlenecks has been effective at removing the most serious deleterious alleles from the populations (Robinson, J. A. et al. 2018; Grossen et al. 2020; Khan et al. 2021; Pérez-Pereira et al. 2022). Our demographic analysis suggests that there have been both historic (∼1900 years ago) and recent (∼170 years ago) bottlenecks in the native eastern black rhino populations. This is consistent with global analyses suggesting that black rhino populations have been persisting in small populations for at least last 2,500 years and have been declining for the last 200,000 years (Moodley et al. 2020). Such bottlenecks are expected to reduce deleterious allele loads but also drive deleterious mutations to fixation and increase homozygosity (Robinson, Jacqueline et al. 2023). Interestingly, the Moru population, which has experienced some gene flow from Ngorongoro (oND and rND cohort), shows lower homozygosity of deleterious mutations than native individuals that had not left the Ngorongoro subpopulation, emphasising the potential benefits of allowing natural movement between populations.

Enrichment analysis of LOF mutations provides a warning that potential fitness consequences should be monitored in wild populations that have introgressed with captive-bred individuals. This is because deleterious mutations can be “hidden” both by heterozygosity in captive populations and because the selection pressures on wild individuals is very different from captive populations, emphasising the role of the environment in expression of inbreeding depression (Keller & Waller 2002). For example, deleterious alleles associated with the myosin and energy-related functions that we identified could potentially affect muscle-related activities, including heart disease, developmental defects or anatomical anomalies (Finno et al. 2009; Finno 2020). This could help to explain the observation that a translocated individual in Grumeti was suspected to have died due to a muscle-related problem (Eblate Mjingo, personal observation). Such health problems might not be apparent in captive populations due to the high nutrition and benign environmental conditions under which they are kept, as well as increased heterozygosity resulting from mixing of individuals from different source populations. However, there also could be impacts of the physical stress of translocating individuals to a novel environment. For example, hemosiderosis, which results from accumulation of iron deposits (hemosiderin) in tissues, was found to be prevalent in captive black rhinos from a UK zoo but also in individuals that had been translocated within Zimbabwe from the wild to managed ranches (Kock et al. 1992). In contrast, high levels of hemosiderin were not observed in free-ranging individuals. Such diseases could also be exacerbated by the diet in captive populations. For example, a serious skin disease (similar to necrolytic dermatitis) was identified in nearly 50% of captive black rhinoceros individuals across the 21 zoo populations that were held in the U.S.A. in the late 1990s (Munson et al. 1998). Since the disease had not been reported in wild individuals, the authors suggested that it could have resulted from metabolic changes due to the rich captive diet. The small size of the native populations could also lead to introduced deleterious alleles reaching fixation rapidly, which may push populations to extinction (Whitlock 2000). Moreover, a further caution comes from the history of the South African captive populations that have been used for translocations to Ndasiata and Grumeti: there are reports of hybridisation between eastern black rhinoceros (*D. b. michaeli*) individuals from Kenya and southern black rhinoceros (*D. bicornis minor*) from Zululand (Hall-Martin 1984). Such admixture between subspecies could have introduced deleterious alleles or contributed to sheltering of the load due to increased genome-wide heterozygosity.

Even though the Serengeti-Mara consists of continuous, unfenced protected areas where movement of individuals with the large home ranges typical of rhinoceros should be possible (Sinclair & Arcese 1995), the intensive protection zone (IPZ) strategy, in which animals are artificially pushed back into specific areas of the landscape where they can be easily monitored, reduces any prospect of natural dispersal between different subpopulations (Fyumagwa & Nyahongo 2010). Thus, the current management strategy would require modifying so that animals are allowed to mix across management boundaries. Previous translocations of eastern black rhinoceros have not considered genetics but our results suggest the potential benefits of capitalizing on the existing variation in the local native populations, rather than relying only on long-distance translocations. This is further emphasized by the observation that the transborder Nyamalumbwa subpopulation, for which mixing is allowed with the Kenyan Maasai Mara subpopulation, shows lower levels of inbreeding than other Serengeti populations. The observational pedigrees could be used to identify unrelated individuals for translocation (Muya et al. 2011), but the approach would be more powerful if combined with an assessment of the genetic load. For example, removing dominant males that have contributed multiple generations of offspring (e.g. in Moru and Ngorongoro) could allow a wider range of individuals to breed, as has been suggested for southern white rhinoceros managed in a metapopulation structure in Botswana (Purisotayo et al. 2019). Genome-wide sequencing data then could be used to model what impacts such a strategy could have on purged deleterious mutations. Nevertheless, our results suggest that sustainable strategies for inbreeding reduction through natural dispersal may be more important than supplementing variation (e.g. increasing heterozygosity) through translocation and reintroduction of captive animals.

Our results showed little effect of management strategies on genetic diversity within cohorts, with a substantial increase in pairwise nucleotide diversity only for individuals whose parents were from captive populations. This is consistent with a recent study using the D-loop of mitochondrial DNA, which suggested that some of the historical maternal diversity in Tanzania had been restored in the populations that included translocated individuals from South Africa or European zoos (Mellya et al. 2023). The study also confirmed that the translocated individuals were from maternal ancestors originally captured from wild populations in Kenya, where many of the lineages still persist. Since Kenyan populations have been found to harbour higher genetic diversity than the Tanzanian populations and there is no fence between the Serengeti (Tanzania) and Maasai Mara (Kenya) management areas, cross-border gene flow could enhance genetic variation. There has already been increased collaboration between rhinoceros management teams in the recent translocation of five eastern black rhinoceros from Ngulia Rhino Sanctuary in Kenya to Ngorongoro in Tanzania for the purpose of increasing diversity, but this was conducted without first obtaining genetic profiles of the translocated rhinos, which would have allowed for prediction of the genetic impacts.

In conclusion, facilitating natural dispersal seems to be the best strategy for managing threatened wild populations like black rhinos, which have been sufficiently bottlenecked to purge out some of the most serious genetic load. Corridors that facilitate animal dispersal have the combined benefits of reducing inbreeding as well as reducing homozygosity of deleterious alleles, thereby reducing the genetic load while maximizing breeding opportunities with unrelated individuals. While translocations from managed game reserves (similar to captive populations) do reduce inbreeding and increase genetic diversity, they risk increasing deleterious allele loads. As the genetic load is sheltered due to potentially benign environments and high heterozygosity in captive populations, there is a risk of future exposure of fitness-reducing deleterious mutations after translocation, unless sustained efforts are made to ensure inbreeding avoidance (Pérez-Pereira et al. 2022). This could be accomplished by changing management practices to allow for natural mixing, which would incur a lower financial cost and improved animal welfare. Alternatively, targeted translocations in each generation could also reduce the risk of inbreeding within the reintroduced population. The power provided by whole genome sequence data offers the opportunity to move away from an original assumption of *ex situ* conservation that supplementation of any genetic variation will reduce extinction risks; instead, we should consider the functional consequences of population mixing on the re-emergence of deleterious alleles, particularly for highly threatened populations (Mable 2018).

## Methods

### Study Area

We focus on the Serengeti-Mara ecosystem on the border of Tanzania and Kenya because the extant populations represent a range of different management scenarios. Black rhinoceros are restricted to five intensive protection zone (IPZ) regions (Figure 1a), where individuals are free-ranging and unfenced but actively monitored by dedicated wardens (Fyumagwa & Nyahongo 2010). This intensive management strategy, along with past translocations, has allowed for population size to increase from a low of 24 in 1995 to 171 in 2021 (Mellya et al. 2023). It also means that there are reliable details on population founders and observational pedigrees, as well as detailed information about reproductive rates and survival of all individuals. We collated these data and constructed pedigrees for each geographic location in our study using the R package visPedigree (https://github.com/luansheng/visPedigree). Individuals that were sequenced in our study are listed with both their names and DNA extraction numbers. Three subpopulations were established by a small number of native founders after a severe bottleneck: 1) **Moru kopjes** (Moru) in the central part of the Serengeti National Park (Serengeti), was founded by two females (M4-Mama Serengeti and M9-Concave) native to the area and one male (R21-Rajabu) that dispersed from the Ngorongoro Crater in 1994 (Supplementary Figure 1); 2) the **Ngorongoro Crater** subpopulation was established by 13 native individuals and two females (mother NG73-Zakhia and calf NG74-Thandi) that had been reintroduced from Addo Elephant National Park (Supplementary Figure 2); 3) **Nyamalumbwa-Maasai Mara** (Nyamalumbwa) in northern Serengeti (a transboundary population between Kenya and Tanzania) was formed by 10 native individuals (on the Serengeti side of the border) (Supplementary Figure 3). Two populations were formed by reintroduced semi wild and captive individuals: 4) **Ndasiata** (Ndasiata), was formed by individuals translocated from Thaba Tholo but there has been subsequent mixing with native individuals who had migrated from Moru (R24-Obayee) and Nyamalumbwa (ND10-Msafiri) (Supplementary Figure 4); and 5) **Ikorongo-Grumeti** (Grumeti), on the western border of the Serengeti, was formed by a young bull (Limpopo) and cow (Laikipia), born at Port Lympne Wild Animal Park. The individuals were translocated to Grumeti Reserve in 2007. Unfortunately, Limpopo was killed in a fight with a bull elephant in 2009. Therefore, in 2018, a young bull from San Diego Zoo Safari Park in the USA (Eric) joined the original female (Laikipia) in the sanctuary but they have not yet produced a calf. Furthermore, in 2019, nine eastern black rhinoceros from South Africa’s Thaba Tholo Game Farm were translocated and released in the wild, comprising of five cows and four bulls (two of whom were calves).

### Sample collection and sequencing

Tissue samples from ears of rhinos were collected opportunistically when the individuals were chemically captured/immobilized for marking, translocation, ear notching operations, fitting of telemetric gadgets, health treatment, or rescue (Supplementary Table 1). The samples were stored in absolute ethanol in 30 ml vials and transported to a laboratory, where they were stored in -20°C until further use. Blood samples were collected for three individuals from Grumeti and stored on FTA cards (Flinders Technology Associates) at 20°C until further use. DNA was extracted using Qiagen blood and tissue DNA extraction kits (Qiagen Inc, Paisley UK).

Libraries for Illumina short-read sequencing were prepared by Novogene, using their in-house DNA Library Prep Set kit. Briefly, genomic DNA was randomly sheared into short fragments which were end repaired, A-tailed and further ligated with Illumina adapters. The fragments with adapters were amplified using PCR, size selected, and purified. The library was quality checked with a Qubit® 2.0 Fluorometer 2010 (Thermo Fisher Scientific, Cambridge), real-time PCR for quantification and a bioanalyzer (Agilent Technologies Inc., Cambridge) for size distribution detection. Libraries were sequenced on Illumina short-read platforms (NovaSeq PE 150) with an aim to obtain at least 30Gbp data per sample.

### Data processing, variant calling and filtering

The raw FASTQ reads obtained from the Illumina platform were end trimmed using default settings of TRIMGALORE for Illumina (https://github.com/FelixKrueger/TrimGalore). The trimmed reads were mapped to the black rhino reference genome assembly (https://www.dnazoo.org/assemblies/Diceros_bicornis) using bwa mem (Li 2013). The mapped reads were sorted and duplicates were marked with samtools (Danecek, Petr et al. 2021). This created the final binary alignment map (BAM) files. The BAM files were then indexed and Strelka2 germline variant caller (Kim et al. 2018) was used for identifying variants. We limited variant calling only to the long chromosomal scaffolds of the rhino genome assembly to avoid biases later when estimating runs of homozygosity (Shukla et al. 2022). This, however, covers more than 90% of the rhino genome. Overall, we obtained 98,736,095 single nucleotide polymorphism (SNP) loci.

The raw variants identified were filtered using vcftools (Danecek, P. et al. 2011). We removed all indel variants. Any base with a PHRED quality score less than 30 was removed and genotypes with quality score less than 30 were set as missing. Any site with a minor allele count less than 3 and deviating from Hardy Weinberg equilibrium with a chi-squared p-value of less than 0.05 were removed. We also removed any individual that had more than 80% missing data. We then removed any loci that were missing in at least 25% of the individuals. We identified sex chromosomes, as described in Armstrong et al. (2021), which were also removed from analyses to maintain consistency in the estimates for males and females. We then estimated the mean sequencing depth at each locus and retained only those loci that had the mid 95 percentile depth of sequencing. Although there was variation among individuals in terms of sequencing depth and retained loci (Supplementary Table 1), overall, we retained 1,649,646 SNP loci after all the filtering.

### Estimating genetic diversity, inbreeding and heterozygosity

Genetic diversity, as represented by pairwise nucleotide diversity (pi), was estimated using the filtered SNP loci using the *--site-pi* option in vcftools (Danecek, P. et al. 2011) for each cohort; to standardise sample sizes, three random samples from each cohort were used.

Levels of inbreeding can be predicted across different historical time periods, based on the length of homozygous tracts spread throughout the genome (runs of homozygosity, ROH), under the assumption that recombination will break up linked variants over time (Broman & Weber 1999). We used BCFtools ROH (Narasimhan et al. 2016) to estimate ROH and then calculate F_ROH_ to measure inbreeding, as described in Armstrong et al. (2021). For each sample we used the ROH option of bcftools and set –G 30 and the allele frequencies were estimated on the “fly with by setting” *–e -*. We then estimated F_ROH_ for each size class using the formula F_ROH_ = length of genome in ROH/total autosomal length.

Overall estimates of heterozygosity of individuals in different cohort categories were defined as the number of heterozygous loci divided by the total loci with data for that individual. We calculated this using RTGtools (https://github.com/RealTimeGenomics/rtg-tools).

### Estimating recent demographic history

We estimated the recent demographic history of eastern black rhinoceros using only the native individuals, by combining the NN, oND and rND cohorts. We used the default parameters in GONE (https://github.com/esrud/GONE) (Santiago et al. 2020) for estimating demographic history. We plotted the harmonic mean of effective population size (Ne) obtained for the first 350 generations, using a generation time of 24 years (Moodley et al. 2020).

### Identifying ancestral alleles, mutation load and homozygosity of deleterious mutations

We defined ancestral alleles as the most common variant present in taxa related to the black rhino. For this, the genome assembly of the northern white rhinoceros (*Ceratotherium simum* genbank assembly: GCA_004027795.1), greater Indian rhinoceros (*Rhinocerous unicornis* genbank assembly: GCA_018403435.1) and Sumatran rhinoceros (*Dicerorhinus sumatrensis* genbank assembly: GCA_014189135.1) were downloaded and used. The data from these three related taxa were mapped to the black rhino genome (https://www.dnazoo.org/assemblies/Diceros_bicornis). From the mapped sequences, the alleles that are most common in the three species were identified as ancestral, as described in Khan et al. (2021).

Under the assumption that derived alleles are expected to more often be deleterious than ancestral variants (Khan 2023), we used the approach described in Khan et al. (2021) to quantify the genetic load. Briefly, the initial set of variants identified from Strelka2 (Kim et al. 2018) were filtered to remove indels, bases below PHRED quality 30 and genotypes below a score of 30. Individuals with more than 80% missing data were removed. Then, we removed any loci missing in at least 25% individuals. Loci with minor allele count of at least 1 and falling within the mean depth across all loci in the mid 95^th^ percentile were retained. Loci with F_IS_ values of - 0.5>F_IS_>0.95 were retained and sex chromosome scaffolds were removed.

The relative genetic load (i.e. relative fitness reduction due to accumulation of deleterious mutations) of pairs of donor and recipient populations was predicted for each cohort type based on both missense (i.e. amino acid changes that retain the function of a protein) and more serious loss of function (LOF) mutations (Robinson, J. A. et al. 2019; Saremi et al. 2019; Khan et al. 2021). The SNPs obtained were annotated using the black rhinoceros assembly annotation files using ensembl VEP (McLaren et al. 2016). Loci with missense mutations or LOF, as defined in (Xue et al. 2015), were identified and the derived allele at these loci was classified as deleterious. All intergenic regions were classified as neutral sites. We randomly selected three individuals from each cohort for estimating mutation loads for all pairs of cohorts to control for the differences in sample size. The R_XY_ method, described by Do et al. (2015) and as implemented by Xue et al. (2015), was used to estimate load. Standard deviations were obtained by 100 rounds of bootstrap. Since most deleterious alleles are recessive in nature (Robinson, Jacqueline et al. 2023) and deleterious only when homozygous, to infer the realised genetic load, we estimated the number of loci with homozygous derived LOF and derived missense mutations and then divided by the total number of loci hosting derived LOF and missense mutations.

### Enrichment analysis

In order to determine what types of processes might be associated with the most serious mutations, WebGestalt (Liao et al. 2019) (http://www.webgestalt.org) was used for gene enrichment analysis of the LOF alleles. *Homo sapiens* was set as the organism of interest, the Over-Representation Analysis was chosen as the method of interest and gene ontology Biological Process, Cellular Component and Molecular Function was the functional database that was chosen. The black rhinoceros genome annotation was uploaded as the reference set. We then uploaded the LOF mutation-containing gene list to run the analysis, with default parameters.

## Acknowledgements

We thank the following institutions for making tissue samples available for this study: Tanzania National Parks (TANAPA), Tanzania Wildlife Authority (TAWA), Ngorongoro Conservation Authority (NCA) and the Kenyan Wildlife Service (KWS). We also thank the Tanzania Wildlife Research Institute and Tanzania Commission for Science and Technology (COSTECH; 2021-449-NA-2021-150) for granting permits to conduct the study. RVKM was supported by grants from Zurich Zoo, the Paul Tudor Jones Family Trust, and a National Geographics Society early career grant (EC-51221R-18). AK was supported by a grant from the Leverhulme Trust (RPG-2020-081). For the whole genome sequencing, we worked in collaboration with Novogene, who provided exceptional support during the optimization phases. We sincerely thank Umer Zeeshan Ijaz, who generously provided access to his server, even though this project was not included in our contributions to his cluster. We also greatly benefited from useful discussions with Mike Bruford, prior to his untimely passing; his insights and generosity will be greatly missed by the conservation genetics community.

## Notes

### Competing Interest Statement

The authors have declared no competing interest.

