## Supplementary Figures for "Natural dispersal is better than translocation for reducing risks of inbreeding depression in eastern black rhinoceros (*Diceros bicornis michaeli)*"

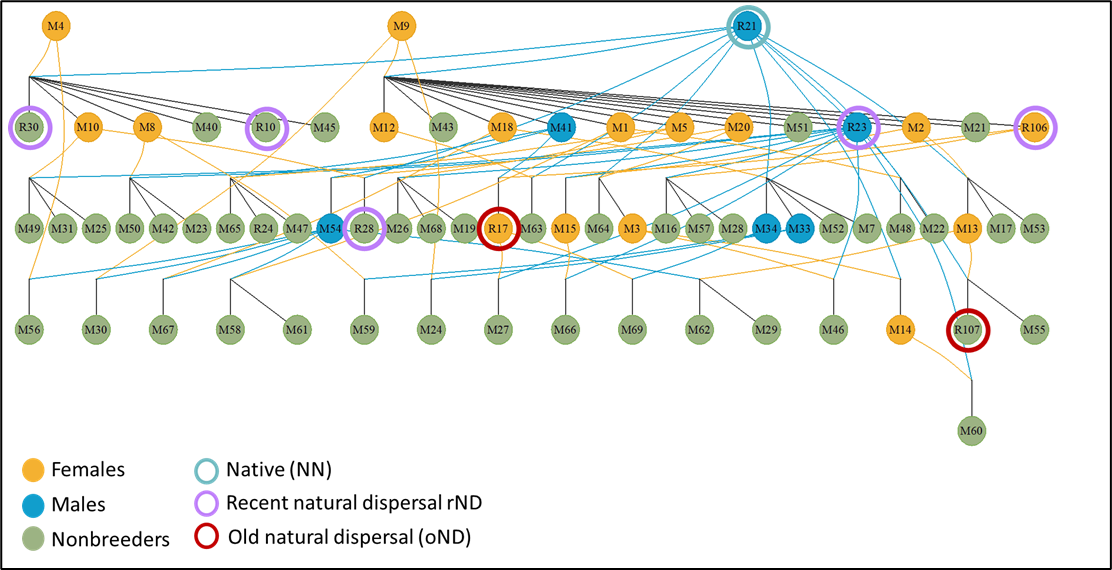

**Supplementary Figure 1.** **Observational pedigree of the Moru subpopulation.** The population was founded by two females native to Moru (M4 and M9) and one male (R21), which had migrated by natural dispersal from Ngorongoro. The dark sky-blue circles denote males, dark golden yellow circles females, and dark olive-green circles nonbreeders, which are either sub-adults or adults without offspring. Ancestors are at the top, descendants in the subsequent rows and parents and offspring are connected by lines. The individuals we sampled for this study are circled with the colour of their cohort type (teal = native individuals whose parents have not dispersed from another population, NN; purple = individuals whose parent dispersed between native populations, rND; red = individuals whose grandparent or great-grandparent dispersed between native populations, oND). The other cohort categories (TC and AD) are not represented in this subpopulation.

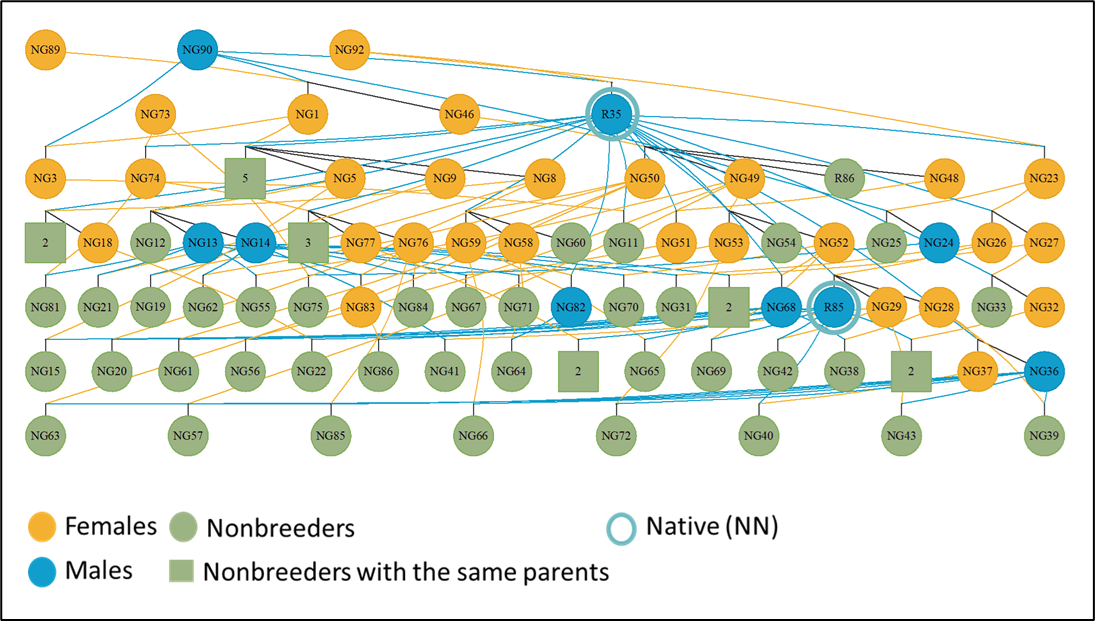

**Supplementary Figure 2. Observational pedigree of the Ngorongoro subpopulation**. The population was formed by 13 native rhinos and 2 females (mother NG73-Zakhia and daughter NG74-Thandi) that had been reintroduced from Addo National Park. The daughter went on to produce six offspring (NG80, NG81,NG79, NG78, NG77, NG76). For this study, we only sequenced individuals that were classified as native, non dispersed (NN; circled in teal); this included an individual who had moved to Moru and so is listed on that pedigree (R21-Rajabu), because the cohort classification is based on the original location of the parents but the pedigrees are based on current location. Given the large number of individuals in this subpopulation, full-sibs who have not left offspring (nonbreeders with the same parents) have been collapsed into green squares, with the number indicating the family size.

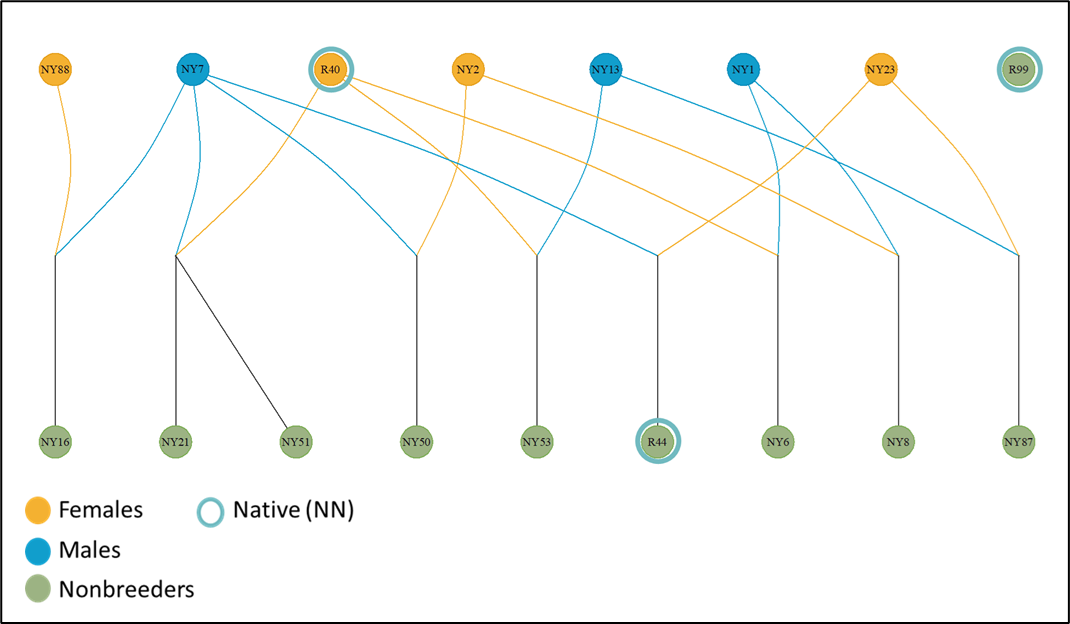

**Supplementary Figure 3.**  **Observational pedigree of the Nyamalumbwa subpopulation**, which is a transborder population between Tanzania (Serengeti National Park) and Kenya (Maasai Mara) and has been formed exclusively by native eastern black rhinoceros. The individuals we sampled for this study (R44, R40 and R99) were all classified as NN (circled in teal).

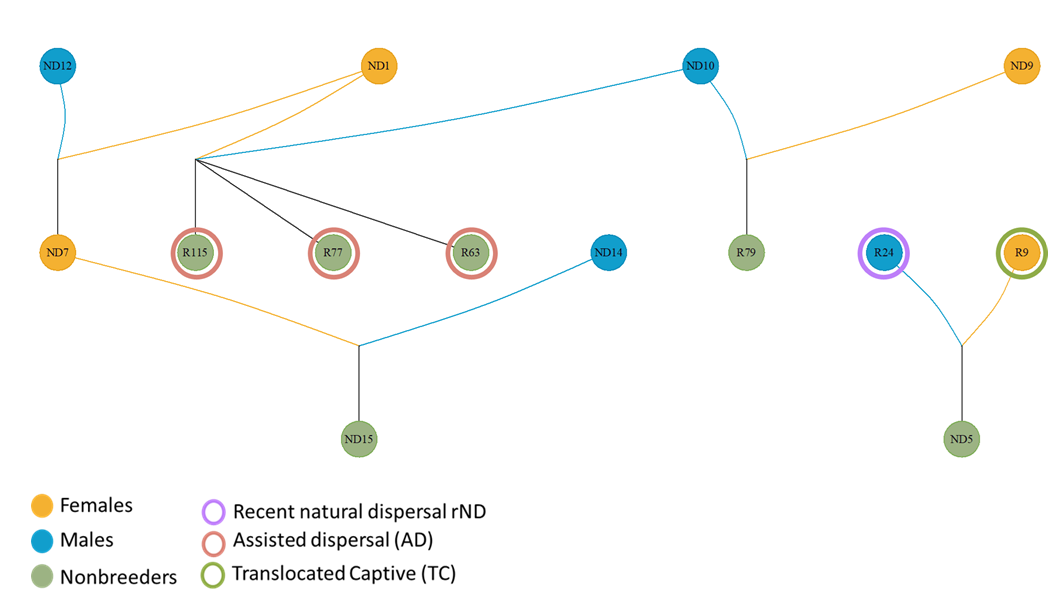

**Supplementary Figure 4.** **Observational pedigree of the Ndasiata subpopulation**. The population was formed by the reintroduction of three captive females (R9, ND9, ND1) from Thaba Tholo Sanctuary in South Africa. Subsequently, a native male from Moru (R24-Olbaye), which was himself the offspring of an individual who had migrated from Ngorongoro and so classified as rND in our sequencing) and Nyamalumbwa (ND10) migrated to the area and bred with the captive females. We sequenced three offspring (R115, R77 and R63) resulting from mating between translocated and native individuals and so classified as assisted dispersal (AD; circled in orange). We also sequenced one of the founders (R9) from Thaba Tholo (classified as translocated captive, TC; circled in green); since there was only one individual of this type from Ndasiata, she was grouped with the Grumeti population for the cohort analyses.

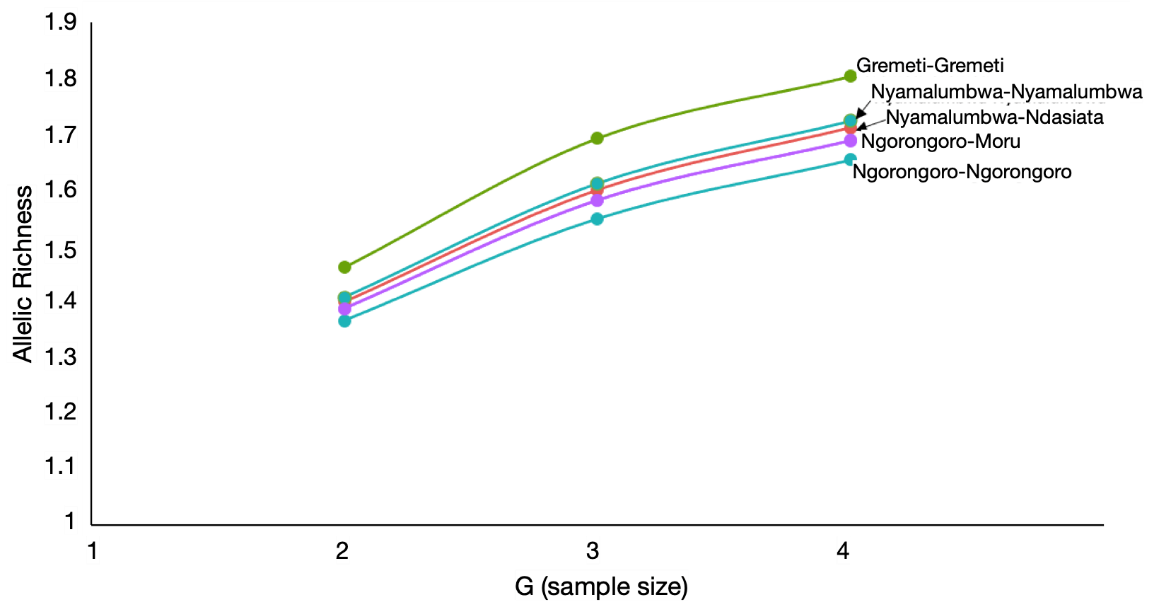

**Supplementary Figure 5.** Allelic richness in each cohort type. The Y axis represents the average number of distinct alleles per locus and the x axis represents the sample size taken from each cohort during a round of rarefaction. oND and rND overlap completely and so are represented as Ngorongoro-Moru.

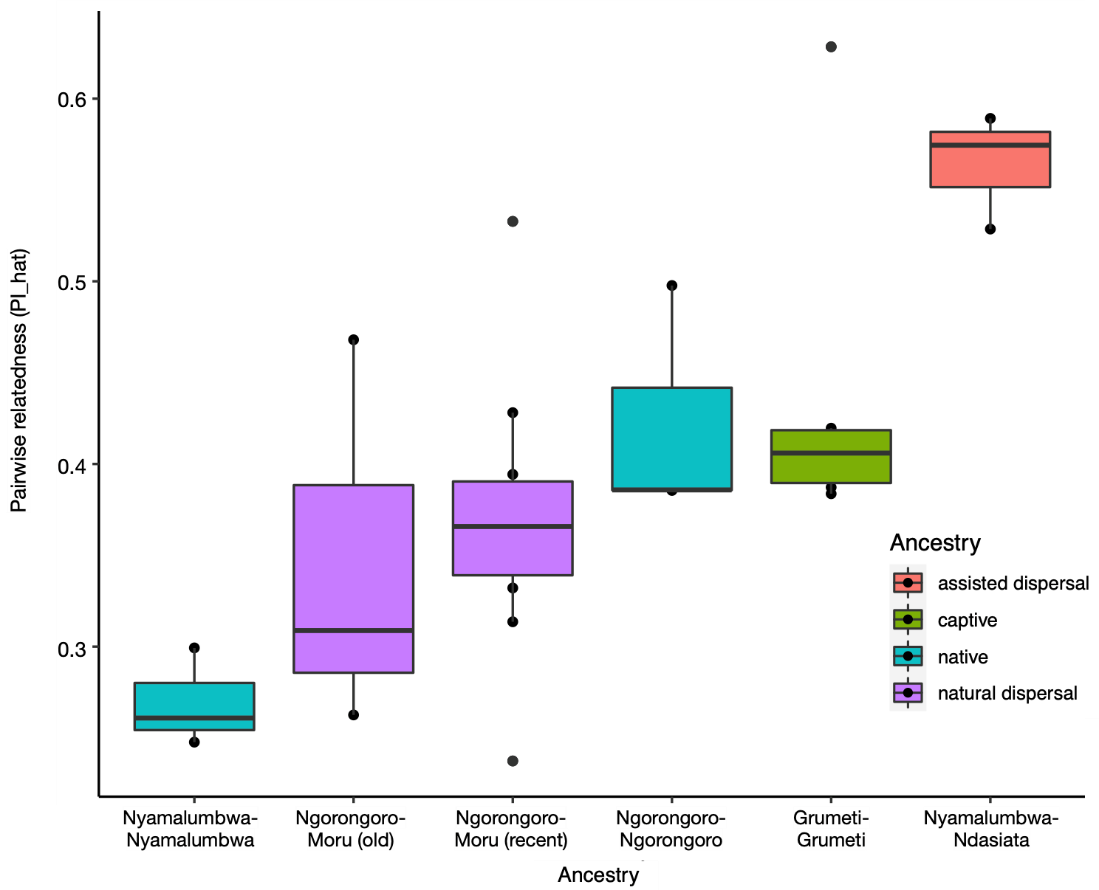

**Supplementary Figure 6.** Pairwise relatedness among individuals in each cohort

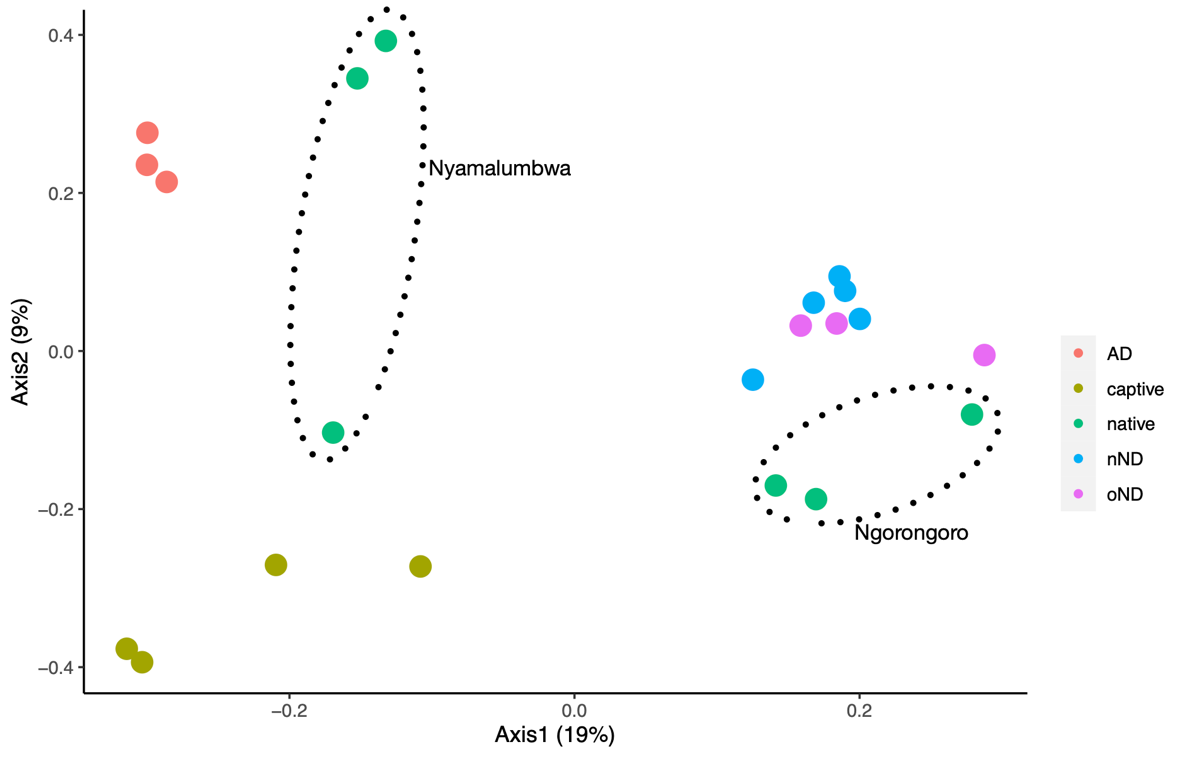

**Supplementary Figure 7.** Principal components analysis (PCA) from single nucleotide polymorphisms (SNPs) of all individuals in the dataset. The individuals from the two NN cohorts have been circled.

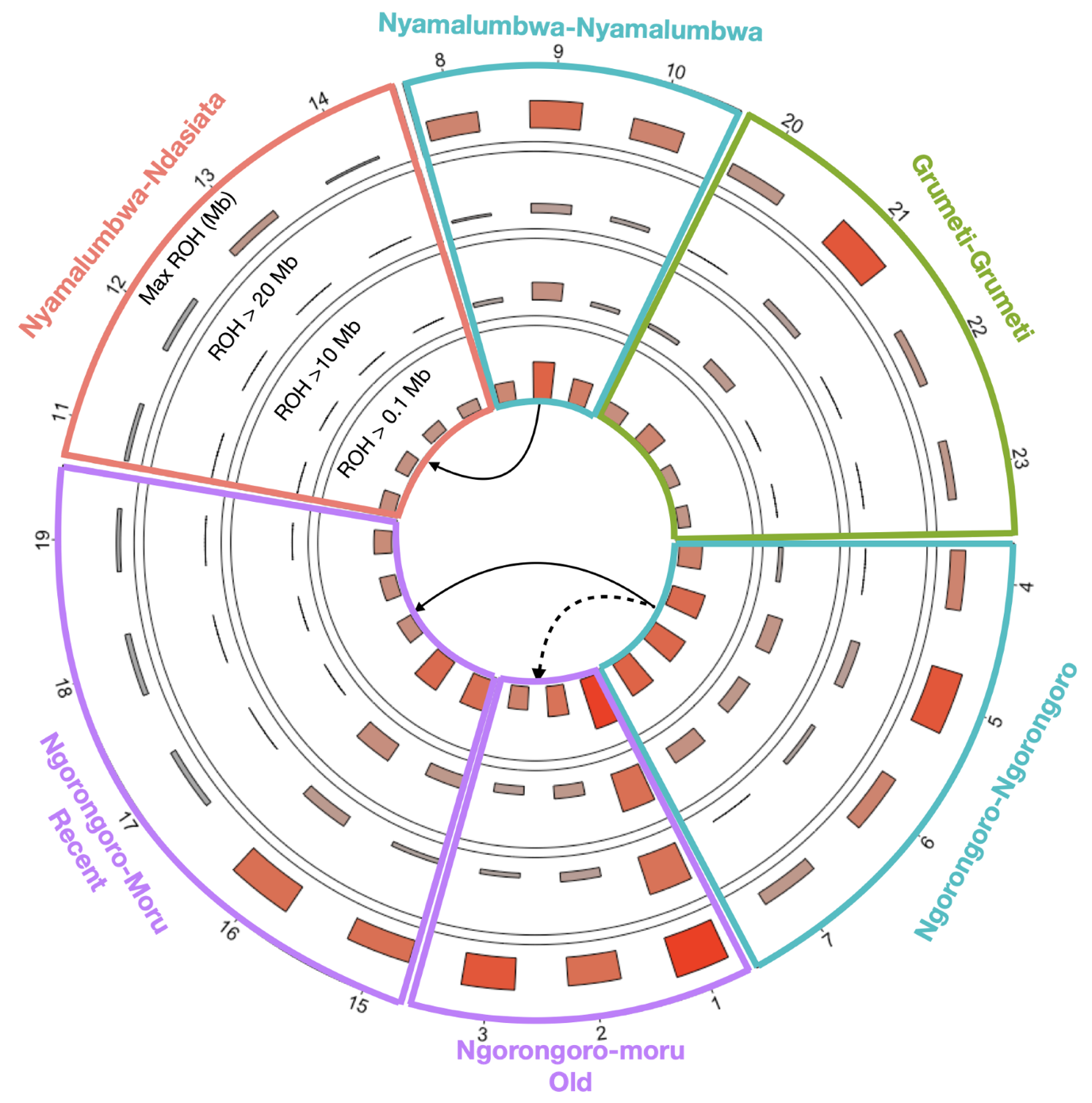

**Supplementary Figure 8**. Runs of homozygosity of various lengths for each cohort, reflecting inbreeding over different historical time periods. A critical assumption is that blocks of homozygosity will be broken down by recombination over time, so the longer tracts reflect more recent inbreeding. The arrows indicate dispersal of individuals between native populations. The numbering of individuals is arbitrary but clustered by population and ancestry cohort. Note that individuals classified as assisted dispersal (Nyamalumbwa-Ndasiata) show the lowest levels of inbreeding, across all time periods. The translocated captive individuals (Grumeti-Grumeti) show relatively low historical inbreeding but individual 21 showed more substantive inbreeding, comparable to some of the individuals in the native populations (Ngorongoro-Ngorongoro). Note that individuals that had recently migrated naturally between Ngorongoro and Moru showed lower levels of inbreeding than earlier generations that had moved.

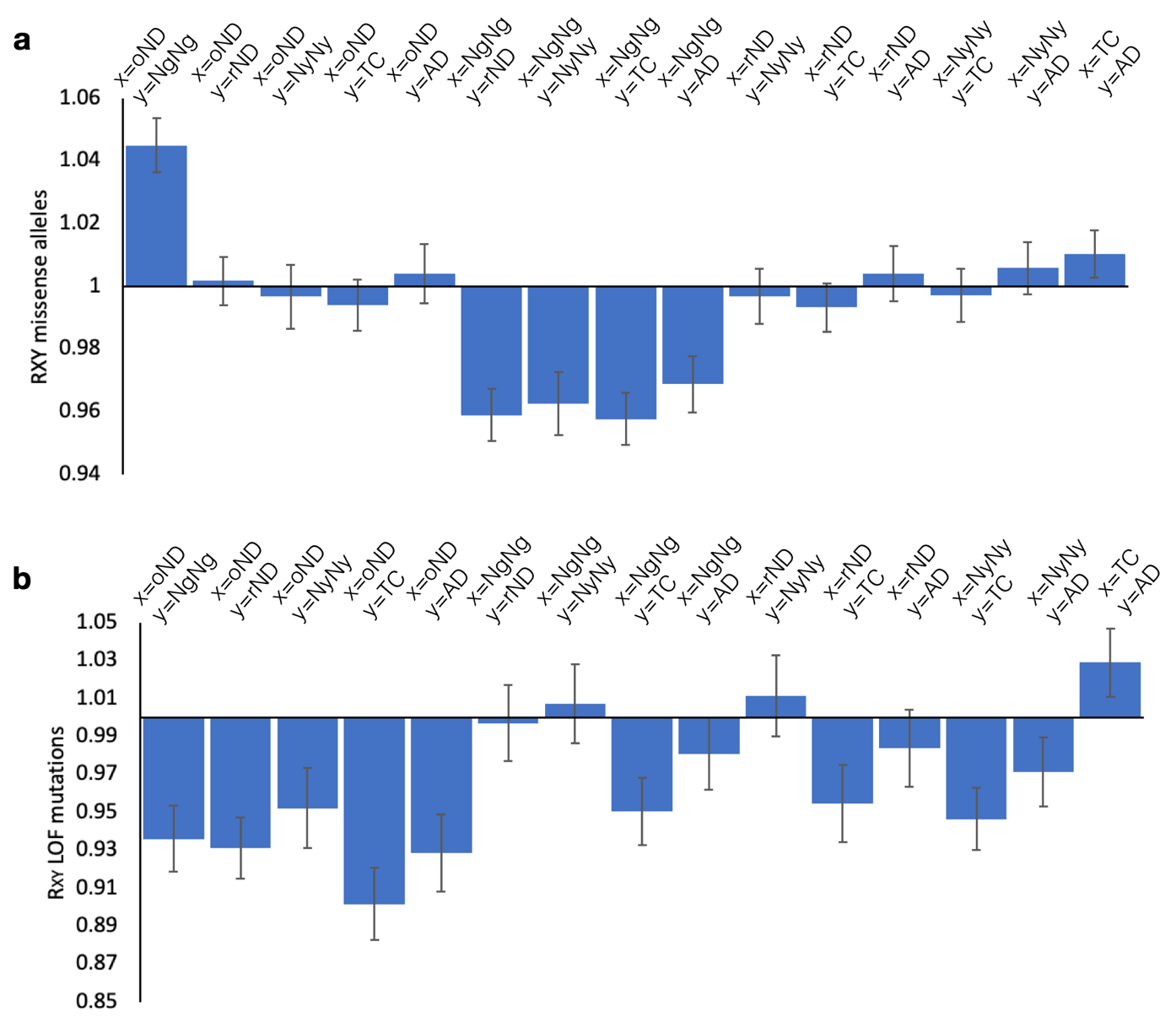

**Supplementary Figure 9**. Deleterious allele loads (Rxy), measured as that in the first cohort “x “with respect to the second cohort “y” listed between all populations. (a) missense allele load and (b) loss of function allele load. Cohort ancestry is indicated as follows: NgNg = Ngorongoro-Ngorongoro; NyNy = Nyamalumbwa-Nyamalumbwa; AD = Nyamalumbwa-Ndasiata; TC = translocated captive (including individuals from Grumeti and Ndasiata); rND and oND are recent and old natural dispersal from Ngorongoro to Moru.

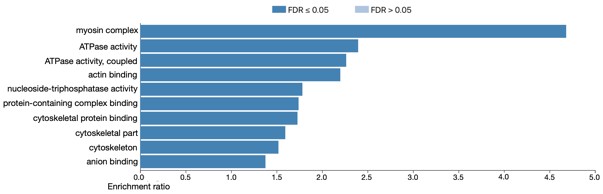

**Supplementary Figure 10.** Enrichment analysis for LOF mutations. Several myosin complex related genes have LOF mutations.

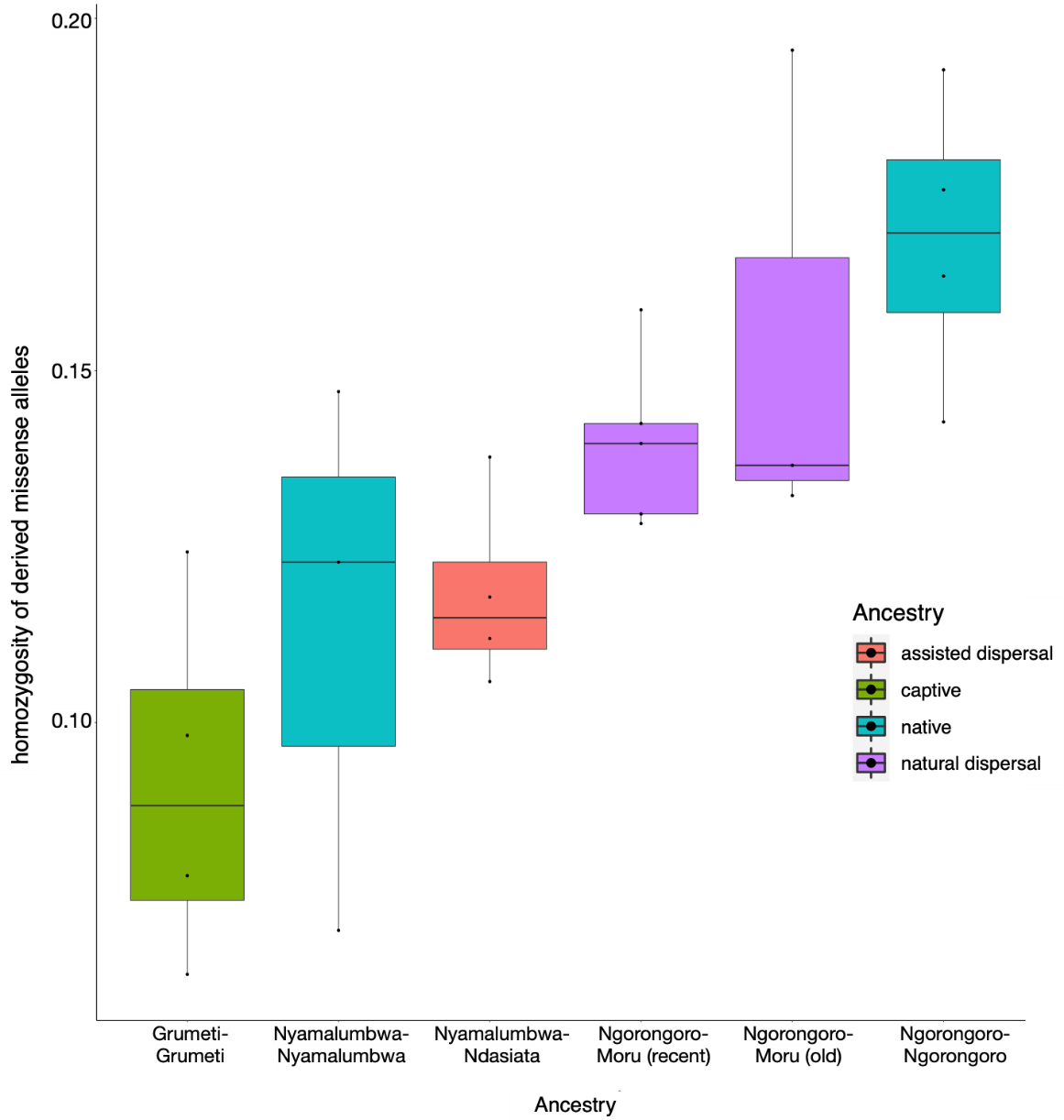

**Supplementary Figure 11.** Homozygosity of derived missense alleles per cohort. Note that the native Ngorongoro populations have the highest homozygosity but they had a lower genetic load than the other categories, whereas the reverse was true for the captive individuals translocated to Grumeti.
